# Boundaries in Spatial Cognition: Looking like a Boundary is More Important than Being a Boundary

**DOI:** 10.1101/391037

**Authors:** James Negen, Angela Sandri, Sang Ah Lee, Marko Nardini

## Abstract

Large walls and other typical boundaries strongly influence neural activity related to navigation and the representations of spatial layouts. They are also major aids to reliable navigation behavior in young children and non-human animals. Is this because they are physical boundaries (barriers to movement), or because they present certain visual features, such as visually extended 3D surfaces? Here, these two factors were dissociated by using immersive virtual reality and real boundaries. Eighty adults recalled target locations in one of four environments: *plywood*, where a virtual wall coincided with a large piece of real plywood; *pass through*, where the virtual wall coincided with empty space and participants could pass through it; *pass over*, where the virtual wall was projected downward to be visible underneath a transparent floor; and *cones*, where the walls were replaced with traffic cones. One condition had features that were boundaries and looked like boundaries (*plywood*); two had features that were not boundaries but looked like boundaries (*pass over/through*); and one had features that were not boundaries and did not look like boundaries (*cones*). The precision and bias of responses changed only as a function of looking like a boundary. This suggests that variations in spatial coding are more closely linked to the visual properties of environmental layouts than to whether they contain physical boundaries (barriers to movement).

---

Spatial memory and navigation are among the most important and widely-shared cognitive tasks performed by humans and other organisms. In the study of spatial cognition, there has been wide interest in differences that appear when presenting typical boundaries (e.g., walls) versus non-boundaries (also called landmarks, beacons, or signs) in the environment. These differences are evident in a wide variety of experimental measures and settings, including in learning (e.g. Austen & McGregor, 2014; Doeller & Burgess, 2008; Pearce, 2009), neural representations (Lever, Burton, Jeewajee, O’Keefe, & Burgess, 2009; Solstad, Boccara, Kropff, Moser, & Moser, 2008) and systematic biases (e.g. Batty, Spetch, & Parent, 2010; Kosslyn, Pick, & Fariello, 1974; Newcombe & Liben, 1982). These differences are not only reported in studies with adult humans, but also young children (Cheng, Huttenlocher, & Newcombe, 2013; Lee, 2017), chickens (Lee, Spelke, & Vallortigara, 2012), zebrafish (Lee, Ferrari, Vallortigara, & Sovrano, 2015), and rodents (Cheng, 1986). These findings lend insight into the mechanisms of spatial cognition; they suggest that boundaries in natural scenes play a crucial role in how we and other animals represent and navigate the world around us (Lee, 2017).

Within this field, a central question remains unresolved. Why do the typical boundaries in natural scenes affect spatial cognition differently from the non-boundaries in natural scenes? Imagine a wall versus a traffic cone. Do the walls affect spatial cognition differently because they *are* boundaries (that physically block movement) or because they *look like* boundaries? In other words, is impeding navigation a relevant predictor or are these effects based solely on visual appearance?

**Table 1.**
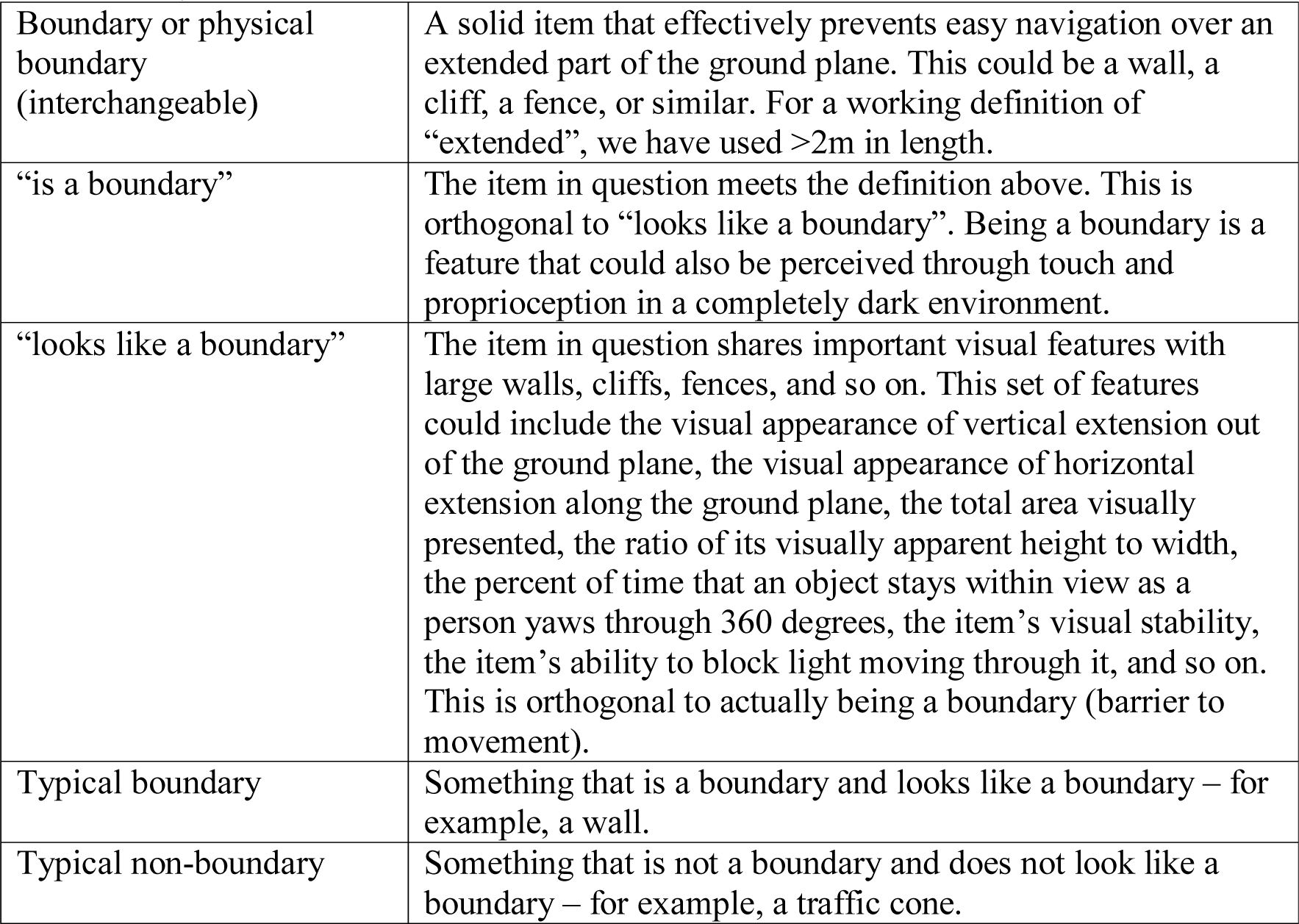
Key Definitions.

This has been a major point of debate for over 30 years. Some articles emphasize the role of a (physical) boundary as an obstacle to navigation, explicitly rejecting a theory based on how they look (Doeller & Burgess, 2008; Doeller, King, & Burgess, 2008). Some articles take the same position implicitly, only describing the differences in terms of how the items impact available navigation and not in terms of how they look (Cheng, 1986; Gallistel, 1990, 2017; Hermer & Spelke, 1994). Some articles conclude that being a boundary is not important, but that visual differences can have a large impact, especially how visually salient the items in the environment are (Austen & McGregor, 2014; Buckley, Smith, & Haselgrove, 2015; Kosaki, Austen, & McGregor, 2013; Mou & Zhou, 2013; Pearce, 2009) and whether they present a visually extended 3D surface (Lee, 2017; Lee, Shusterman, & Spelke, 2006; Lee & Spelke, 2010). The mechanisms underlying spatial cognition in natural environments cannot be understood fully until this is resolved.

We need to separate *being a boundary*, which means impeding navigation across an extended part of the ground plane, from *looking like a boundary*, which means sharing important visual features with large walls. From first principles of experimental design, it is clear that we need to be able to make two comparisons to resolve this question definitively. First, to see if being a (physical) boundary has a specific effect, we need two conditions that look exactly the same but differ in whether the scene contains boundaries. Second, to see if looking like a boundary has an effect, we need two conditions that impede navigation in the same way but differ in whether they look like a boundary.

The gap in the current literature is that this first comparison is not available; there is no published test where the (physical) boundary and non-boundary look exactly the same. However, it has been established that looking like a boundary (e.g., visually presenting a vertically extended opaque surface) has reliable effects on spatial cognition, even when the presented items all impede movement in the exact same way (Gianni, De Zorzi, & Lee, 2018; Kosslyn et al., 1974; Newcombe & Liben, 1982). To be specific, adult participants tend to rank the distance between two targets as longer if a straight line between them crosses an opaque boundary, but not a transparent boundary of the exact same size and shape (Kosslyn et al., 1974; Newcombe & Liben, 1982). Further, young children can reorient to an opaque rectangular boundary but not a transparent rectangular boundary (Gianni et al., 2018). There are also patterns of brain activation that are selectively associated with images of boundaries, even when the participant is not given any opportunity to move or navigate (Park, Brady, Greene, & Oliva, 2011; *see also* Julian, Ryan & Epstein, 2016; Konkle & Oliva, 2012). This answers half of our central question, leaving open whether there is (a) only an effect of looking like a boundary or (b) both an effect of looking like a boundary and an effect of being a boundary.

Another previous study came to a closely-related conclusion (McNamara, 1986). This study included comparisons of spatial cognition in two environments with strings on the ground. In one condition, participants were allowed to move freely over the strings. In the other, they were instructed not to move over the strings. We would refer to this as a *social boundary*. No major difference was found between conditions; the pattern of results was similar in terms of recognition priming (if two objects were near each other or in the same region as each other during the study phase, presenting one during the test phase leads to faster recognition of the other on the next trial), direction judgements (which direction from one item to another), and distance estimation (Euclidian distance from one item to another). Since the strings and instructions were not physically capable of impeding navigation, they were not boundaries under the current definition. It remains unknown if a (physical) boundary effect exists.

The present study employed four conditions, between-subjects, in an immersive virtual reality setting (a simulation traversed by actual movement, not with a controller; Figure 1, Table 2) to fill this gap and to further test for a purely visual effect. The conditions varied both in terms of their virtual environment and in terms of the real space that this was projected onto. The first condition, *plywood*, presented items that are (physical) boundaries and also look like boundaries. These were large blue walls that visually presented an extended surface and were co-located with a large piece of real plywood. Participants touched the plywood to establish that it actually impeded movement. The first comparison condition, *pass through*, was visually identical, but participants established by touch that they could pass through the walls (i.e., there was no plywood). The items were not boundaries but did look like boundaries. The second comparison, *pass over*, used the exact same wall-like items but placed them under the translucent floor, coinciding with the ground plane at their top edge. This means that vision alone could confirm that the items in the environment were not boundaries; an item cannot block movement if it is beneath the floor. To test for a purely visual effect, the fourth condition, *cones*, presented a set of virtual traffic cones that also were not boundaries and did not look like boundaries. Participants directly experienced which conditions contained boundaries and which did not by touching the walls in *plywood* and moving through/over them in the others. The task was to remember the position of three target locations and indicate these again after a change of viewpoint. To measure the extent to which spatial coding was similar or different across these environments, we looked for differences in patterns of recall error – specifically, the bias and precision of responses.

**Figure 1:**
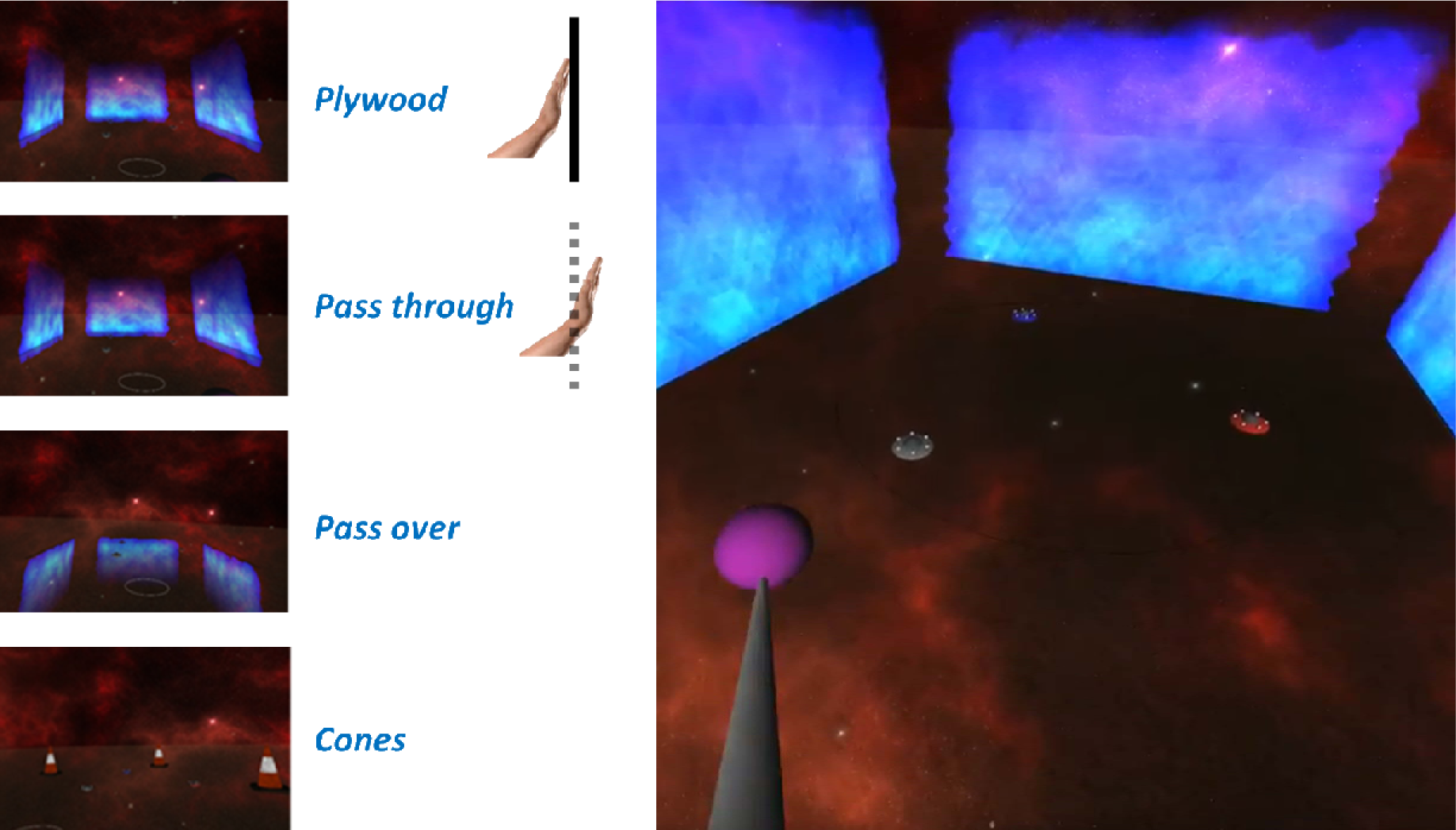
Conditions. The experiment had four different conditions. Participants were immersed in one of three unique virtual reality environments. The *plywood* and *pass through* conditions used the same virtual environment, making them visually identical in every detail. For *pass over*, the walls were under the floor, but the walls themselves joined the floor at the same locations and otherwise looked identical. The *cones* condition replaced the walls with traffic cones. In terms of containing boundaries, only the walls in the *plywood* condition gave any resistance when participants attempted to pass through them. These conditions allow us to dissociate whether the presented items are boundaries (*plywood* contains boundaries, but *pass through/over* and *cones* do not) and whether the presented items look like boundaries (the cones in *cones* do not, but the walls in *plywood* and *pass through/over* do so).

**Table 2.** Reference for the key features of the different conditions.

| <u>Condition</u> | <u>Is a Boundary</u> | <u>Looks Like a Boundary</u> |
| --- | --- | --- |
| Plywood | Yes | Yes |
| Pass Through | No | Yes |
| Pass Over | No | Yes |
| Cones | No | No |

Some previous studies have attempted to understand a special role for boundaries through an overshadowing paradigm (Doeller & Burgess, 2008), but this has failed to provide a consistent boundary versus non-boundary effect (Austen & McGregor, 2014; Buckley et al., 2015; Kosaki et al., 2013; Mou & Zhou, 2013; Pearce, 2009). Further, any approach that only examines accuracy will fail to capture important effects on bias and precision (Huttenlocher, Hedges, & Duncan, 1991; Kosslyn et al., 1974; Newcombe & Liben, 1982). Our approach is to focus on the more basic phenomenon and examine how target locations are encoded and recalled in different environments. This makes the present study similar to the previous social boundary study (McNamara, 1986) and to previous studies with young children (Cheng et al., 2013; Gouteux, Vauclair, & Thinus-Blanc, 2001; Lee, 2017; Lee et al., 2006; Lee, Winkler-Rhoades, & Spelke, 2012). However, instead of expecting a categorical difference (young children can perform above chance in some environments, but not others), we expect different environments to cause graded differences in the bias and precision of responses. This is also why it is important to include the *cones* condition: a difference between *plywood* and *cones* can confirm that the task and analysis are sensitive to the difference between a typical boundary (*plywood*) and a typical non-boundary (*cones*).

Based on previous work (Gianni et al., 2018; Kosslyn et al., 1974; Newcombe & Liben, 1982), we expected to find a difference in terms of *cones* versus *pass through* and *pass over* (i.e. a purely visual difference). The main hypothesis was a difference in performance between *plywood* versus *pass through* or *pass over.* This would suggest that there are both (a) effects of looking like a boundary and also (b) pure effects of being a (physical) boundary, independent of visual appearance. The alternative hypothesis is that there will not be a difference between *plywood* versus *pass through* and *pass over*. This would suggest that there is only an effect of looking like a boundary. This would in turn indicate that the effects of typical boundaries in natural environments are driven by their typical visual aspects rather than directly by their physical limitations on navigable space.

## Method

### Summary

Participants were given a spatial memory and navigation task in one of four conditions through the use of immersive virtual reality (Figures 1 and 2). Participants were shown three target locations (marked by small spaceships) and then ‘teleported’ to a new viewpoint within the arena and asked to either point to the spaceships (spatial memory) or walk over to their locations (navigation) in order. The conditions differed in how they looked and if they contained boundaries.

**Figure 2:**
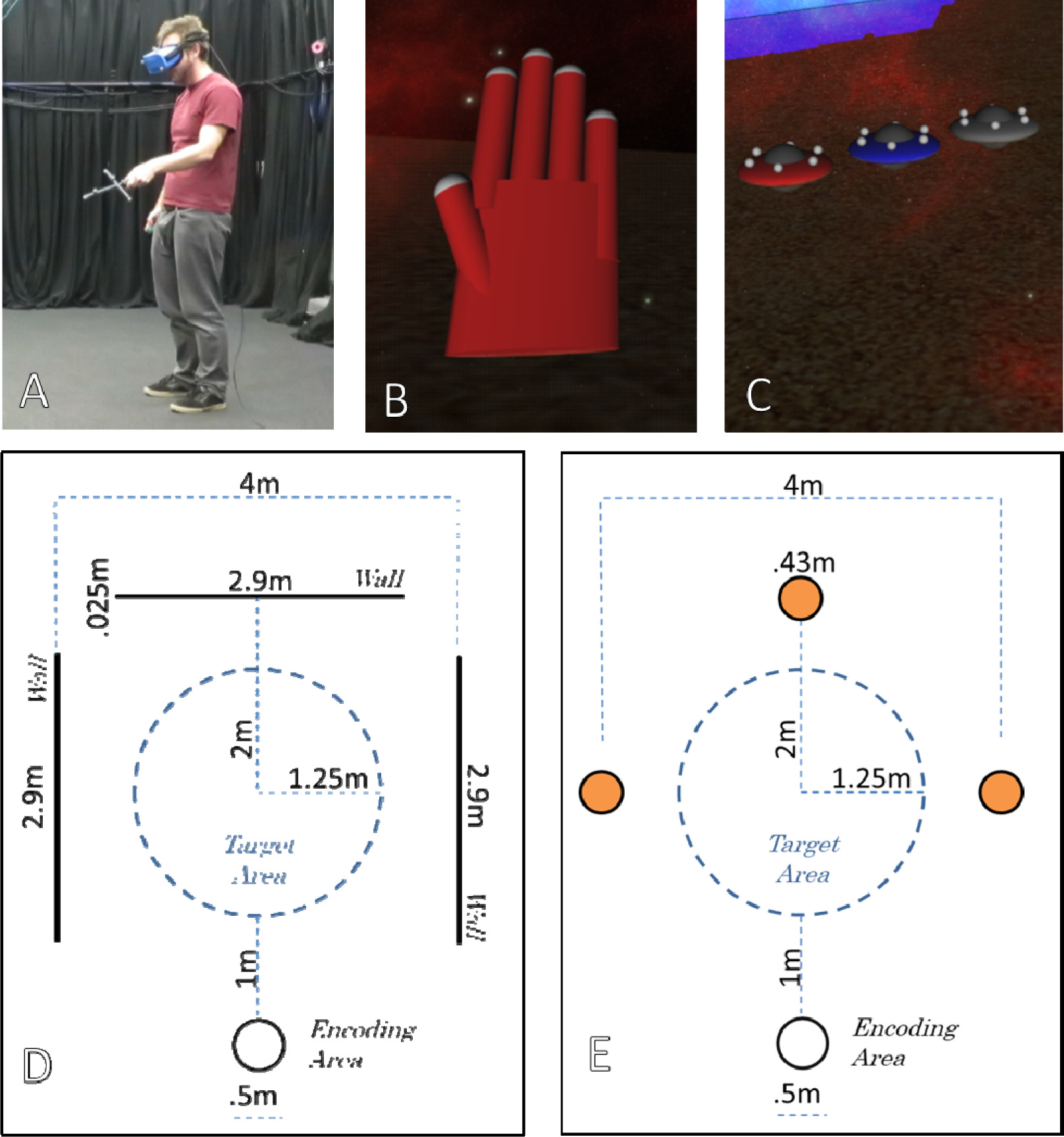
Task Design. (A) A photograph of a person participating in the experiment. (B) The red ‘robot hand’ that moved with the right hand of participants in the virtual world. Before data collection, participants used this to experience what items they could or could not move through. (C) The targets, in a line here to see easily. (D) An overhead diagram of the *plywood, pass through*, and *pass over* conditions. (E) An overhead diagram of the *cones* condition. In short, participants were asked to stand in the Encoding Area and watch the three targets appear in the target area and disappear in order. They were then ‘teleported’ (screen faded to black, ‘camera’ moved, screen came back up) to a point on the edge of the target area. Participants indicated where they remembered the three targets in order.

### Participants

Eighty undergraduates were recruited to participate. They were given either a credit hour towards a program where students volunteer in each other’s studies or £8. The study was approved by the local Ethics committee and participants gave informed consent. Participants were excluded if they had abnormal vision and could not wear contact lenses. Twenty participants each were randomly assigned to the conditions (see below): *plywood* (10 female, mean age = 23.45 years); *pass through* (10 female, mean age = 22.30 years); *pass over* (10 female, mean age = 26.2 years); and *cones* (10 female, mean age = 21.50 years). This sample size was chosen mainly because it is slightly larger than the number of participants in developmental studies that have found related effects (e.g. Lee & Spelke, 2011). The design also has the added advantage of a much higher number of trials per participant (48 here versus 4 in the cited article).

### Apparatus

Participants wore an Oculus Rift headset (Consumer Version; Menlo Park, CA, USA). The headset was tracked continuously via small reflective markers by a Vicon Bonita motion tracking system (Oxford, UK). They were also given a motion-tracked ‘wand’ for indicating responses, made of PVC cylinders and a screwdriver handle (see Figure 2A), plus a small motion-tracked glove on their right hand. The virtual environment was programmed in WorldViz Vizard 5. With real-time updating of the participant’s and headset’s position from the motion capture markers, participants were able to freely traverse a space of about 5m × 10m, with approximately 4m × 4m used during this study.

#### Virtual environment

The virtual environment for the *plywood, pass over*, and *pass through* conditions featured three blue walls set at right angles (see Figure 1). They were adapted from a fire effect in WorldViz Vizard 5. They were formed of a continuous set of translucent hexagons that moved slowly and drifted randomly. This was done so that there would not be consistent local landmarks along the faces of the walls. To enable the *plywood* condition, the virtual wall to their left was rendered in the same place as a real 4ft × 8ft (approx. 1.2m × 2.4m) piece of plywood which had been mounted with steel brackets to form a stable boundary. To enable the *pass through* condition, the virtual wall to their right corresponded to an empty space in the lab which participants could put their hand through without any resistance. For the *pass over* condition, the walls appeared below the translucent floor but could still clearly be seen. In the *cones* condition, there were instead three traffic cones (dimensions conforming to motorway regulation in the United Kingdom) placed where the centers of the walls were in the other conditions. The cones were 67.5 cm in height, 43 cm across the base, and colored orange and white (see Figure 1).

There was also a skybox rendered at an infinite distance with the appearance of outer space nebulae (i.e. a redundant method for participants to reorient themselves, but without providing distance information). The floor surface was made of a translucent sand texture (20% opacity). Participants also saw a virtual projection of their ‘wand’ and a model of a red ‘robot hand’ where their right hand was (Figure 2B). The participants were asked to hold the wand in their left hand when they were not actively using it, but to use both hands to point as accurately as possible when indicating locations during the main task. The motion capture markers on the right hand were placed on the back side of the hand so that they did not interfere with this.

#### Stimuli

On each trial, the stimulus was a set of three target locations on the floor to remember. This was shown by having three small coloured spaceships (Figure 2C) appear and disappear in sequence at different positions. The ships were red, blue, and grey on their outer hulls, with a grey center and six small white lights on their top sides. They appeared by first being shown as a flat disc and then stretching to their full vertical extent. Then they were removed by flattening again and then disappearing. Each was visible for 1s plus or minus the screen refresh time (1/90^th^ of a second). Spaceship locations were randomly drawn from a uniform circular distribution with a radius of 1.25m around the center of the arena (Figure 1D-E, “Target Area”), constrained only in that they could not overlap with each other.

### Procedure

Participants were not allowed to view the actual testing space before entering. They were fitted with the headset outside and then led in. Each of the three conditions had a pre-testing procedure that drew their attention to the items in the environment at the start of each block. For the *plywood* condition, they walked to the plywood boundary and were asked to feel it with their hands. This was important for establishing that the environment contained a real (physical) boundary. To keep the situation reasonably naturistic, we did not ask participants to experience each wall separately. For the *pass through* condition, they walked over to a virtual wall projected into empty space and were instructed to try moving their hand through it. For the *pass over* condition, they were instructed to walk over the same virtual wall and see what the wall below looked like. For the *cones* condition, they were asked to move their hand through a cone.

There were a total of four blocks of twelve trials. At the beginning of each trial, participants stood inside a small white circle. This location was the same on every trial (Figure 2D-E, “Encoding Area”). From this vantage point, all three walls and the targets could be viewed in the same rendered frame within the headset. When the program registered that the participant was inside the white circle and looking towards the target area, the stimuli were displayed.

The next step was effectively a very short disorientation procedure, giving participants an unpredictable new viewpoint without any useful self-motion information. Participants were ‘teleported’ – the screen faded to black, the virtual camera moved, and then faded up from black – to a place around a 2.5m radius circle in the middle of the three walls (Fig. 2D-E, “Target Area”). The white circle was no longer visible. Their view also rotated; they were facing the center of the target area at encoding and were facing the center of the target area when the screen faded up from black. The rotation amounts were random but within the constraint that each block had six rotations under 90 degrees and six rotations above 90 degrees (180 being the max possible), three to the left and three to the right. This was done to enlarge any potential differences between the *plywood* and other conditions, as typical boundaries have a particularly strong and reliable effect on reorientation in terms of both behaviour (e.g. Hermer & Spelke, 1994) and neural representations (Keinath, Julian, Epstein, & Muzzio, 2017). Crucially, participants did not have additional physical contact with the items after the teleport, creating a more naturalistic use of the boundaries after disorientation.

Participants were asked indicate their response by either pointing with the wand or by walking to the target locations. This was done in four alternating blocks of twelve trials. The response modality for the first block was random. A small inverted cone was rendered on the floor to make it as clear as possible where each pointing or walking response would be recorded. In the pointing case, the inverted cone marked the point where a straight line from the wand intersects the floor. In the walking case, the inverted cone was 25cm in front of the center of their head. The inverted cone’s colour was matched to the target. This was repeated for all three targets, with the inverted cones remaining visible from previous responses. The responses were collected as quickly as participants made them (no additional delay was imposed by the experimenters). The targets then re-appeared and the participants were allowed to see the correct target locations. Participants then returned to the encoding area for the next trial. Breaks were given between blocks as needed. Testing typically took 45 to 60 minutes.

This procedure essentially results in 48 trials, each yielding fourteen figures: the placement of each target and each response along each axis of the ground plane, plus the block number and the rotation amount (the size of the angle formed by the encoding location, the center of the target area, and the placement of the participant at the end of the ‘teleport’). This provides a strong basis for assessing both the bias and the precision for each participant.

## Results

Two trials were lost to technical errors, resulting in 11,514 usable responses along two axes. Before the main analysis of both bias and precision, we examined the raw errors (absolute distance along the ground between target and response) across conditions to provide context. The raw errors were 62 cm on average with a standard deviation of 46 cm and a skewness of 1.58 (median 49 cm). An ANOVA on average raw error was used to examine two factors: being a (physical) boundary and looking like a boundary. This is one contrast for *plywood* versus the other conditions and another contrast for *cones* versus the other conditions (Table 2). Neither the effect of being a boundary nor the effect of looking like a boundary was statistically significant (Figure 3, Table 3). Further, all four 95% confidence intervals overlap. This was not due to a floor effect. We examined the targets and responses along the x-axis (left-right in figures), calculating an unstandardized beta coefficient for each participant. The average coefficient was significantly above zero, t(79) = 31.28, p < .001, mean of 0.75 m/m. On the y-axis (up-down in figures), the same effect was seen, t(79) = 33.97, p < .001, mean of 0.73 m/m. The next analysis involved parsing the raw errors into bias and precision.

**Figure 3:**
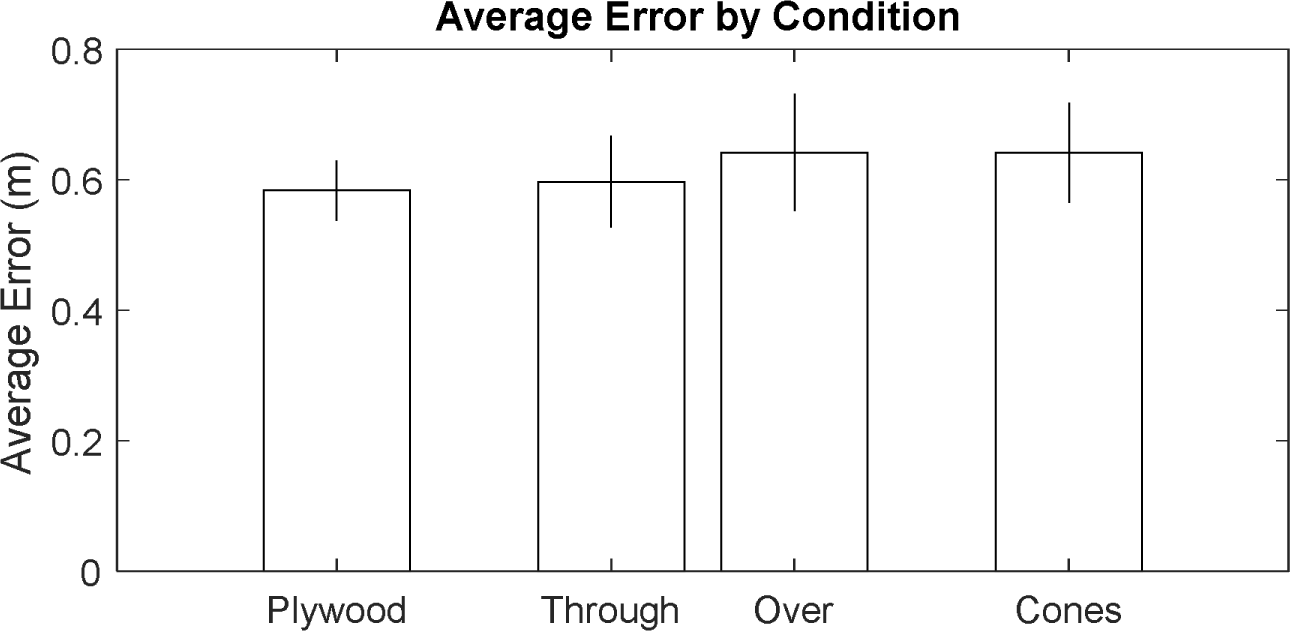
Raw Error by Condition. Raw Error was not significantly different for the different conditions. There was no significant effect of either looking like a boundary (middle two versus far right) or being a boundary (middle two versus far left). Error bars are 95% confidence intervals on the mean of the average error for each participant. All four confidence intervals overlap. For example, an average error of 0.60m is within the interval for all four conditions.

**Table 3.** ANOVA Results.

| Source | DV | SS | d.f. | F | p | $\eta^2$ | $\eta^2$ Upper Bound* |
| --- | --- | --- | --- | --- | --- | --- | --- |
| Is Bound. | Raw Error | 0.02 | 1 | 0.62 | 0.435 | 0.008 | 0.054 |
| Looks like | Raw Error | 0.01 | 1 | 0.24 | 0.625 | 0.003 | 0.067 |
| Error | Raw Error | 2.12 | 77 |  |  |  |  |
| Total | Raw Error | 2.15 | 79 |  |  |  |  |
| Is Bound. | Bias X (Bx) | 0.02 | 1 | 0.28 | 0.598 | 0.004 | 0.068 |
| Looks like | Bias X (Bx) | <0.01 | 1 | 0.01 | 0.940 | <0.001 | 0.048 |
| Error | Bias X (Bx) | 5.27 | 77 |  |  |  |  |
| Total | Bias X (Bx) | 5.30 | 79 |  |  |  |  |
| Is Bound. | Bias Y (By) | 0.02 | 1 | 0.06 | 0.812 | 0.001 | 0.034 |
| **Looks like** | **Bias Y (By)** | **11.09** | **1** | **29.26** | **<.001** | **0.268** | **0.414** |
| Error | Bias Y (By) | 29.19 | 77 |  |  |  |  |
| Total | Bias Y (By) | 41.33 | 79 |  |  |  |  |
| Is Bound. | Bias Weight ($W_B$) | 0.03 | 1 | 0.92 | 0.340 | 0.010 | 0.052 |
| **Looks like** | **Bias Weight ($W_B$)** | **0.26** | **1** | **9.50** | **0.003** | **0.104** | **0.255** |
| Error | Bias Weight ($W_B$) | 2.13 | 77 | | | | |
| Total | Bias Weight ($W_B$) | 2.52 | 79 | | | | |
| Is Bound. | Resid. Error | <0.01 | 1 | <0.01 | 0.991 | <0.001 | 0.042 |
| Looks like | Resid. Error | 0.02 | 1 | 1.09 | 0.300 | 0.014 | 0.098 |
| Error | Resid. Error | 1.23 | 77 |  |  |  |  |
| Total | Resid. Error | 1.25 | 79 |  |  |  |  |

\*Estimated by taking the 95<sup>th</sup> percentile of a bootstrap procedure, where every condition was resampled with replacement 10,000 times and $\eta^2$ was calculated. Significant effects in bold.

We estimated the bias (mean signed error) within several small local areas of the arena, separated by condition, to make flow field charts of the biases (Figure 4). Visual inspection suggests that the bias fields are very similar for the three conditions containing walls that look like boundaries, but that the bias field is not the same for *cones*. The bias field in *cones* converges near the center of the target area, but the bias fields converge further towards the back wall in the other conditions.

**Figure 4:**
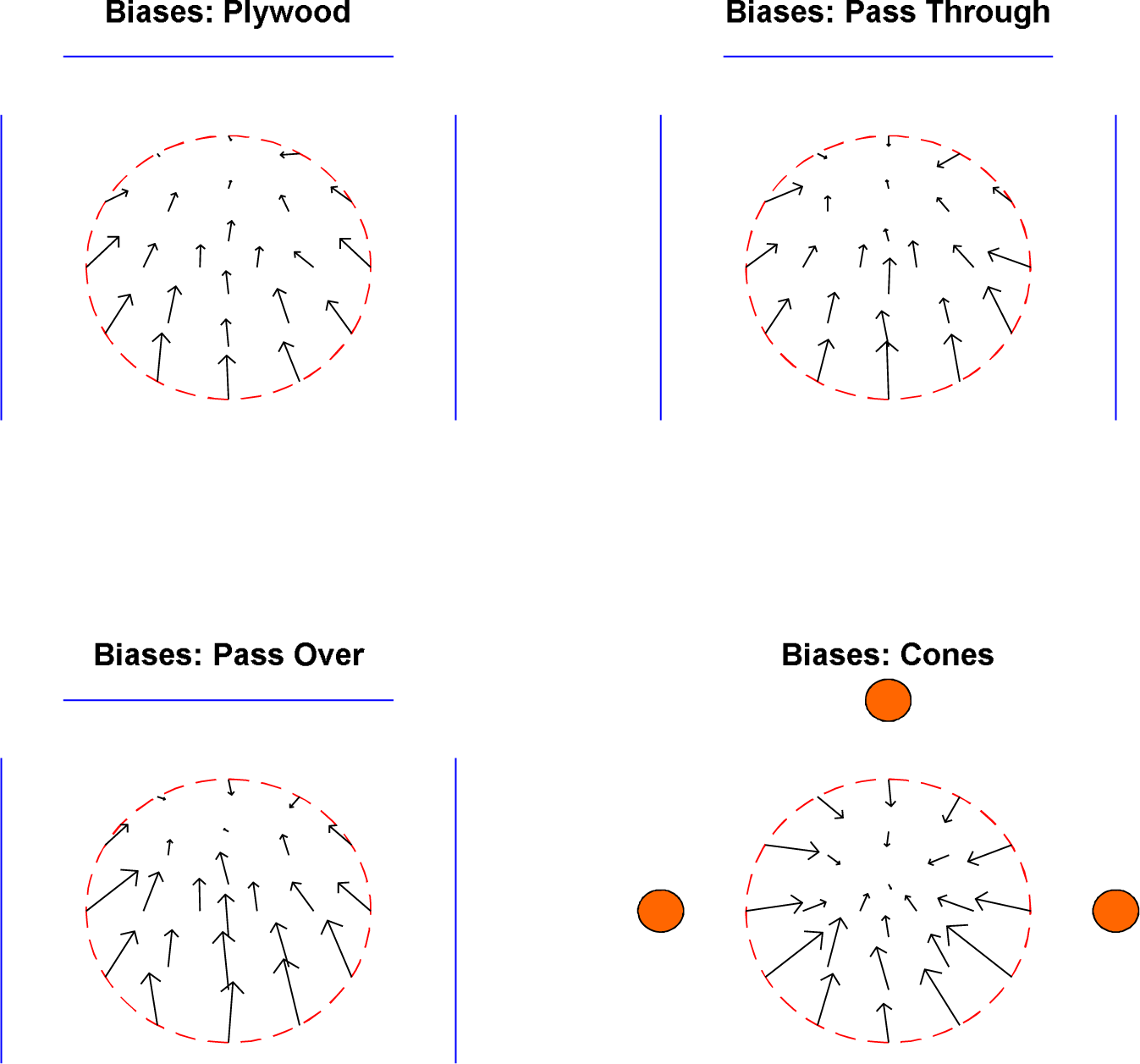
Biases by Condition. The blue lines are the walls and the orange circles are the traffic cones. The red dashed line is the edge of the target area. For each arrow, responses nearest its base had a bias (mean signed error) as indicated by the direction and magnitude of the arrow.

To formally test for differences in bias across conditions, we calculated the bias for each participant. The method assumes that a single point, the bias point, attracts responses towards itself with a certain weight. This is like a simple model of gravity attracting small objects into a gravitational well. To be more specific, we fit three parameters: the bias point’s × value (Bx), the bias point’s y value (By), and the weight given to the bias point (W_B_). To give an example of the relevant calculations, if we present a target at (1, 1) and place the bias point at (0, 0) with a weight of 20%, then we would most expect a response at (0.8, 0.8) since that is 20% of the way towards (0, 0) from (1, 1). Fitting this involved minimizing the squared distance between the observed responses and the expected responses after displacing the targets towards the bias point. This is captured by the equation

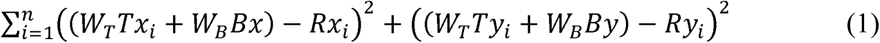

where n is the number of trials, W_T_ is the weight given to the target, W_B_ is the weight given to the bias point, Tx is the target on the x-axis, Ty is the target on the y-axis, Rx is the response on the x-axis, and Ry is the response on the y-axis, with W_T_ + W_B_ = 1, 0 < W_T_ < 1, 0 < W_B_ < 1, −2 < Bx < 2, and −2 < By < 2. The minimum of Equation 1 was found, fitting the three parameters Bx, By, W_B_ (x- and y-positions of the bias point and weight given to the bias). This further allowed us to calculate the residual error as a fourth outcome, which captures the parsed precision. To continue using the example, if we observed a response at (0.7, 0.8), that would be a residual error of 0.1, since we most expected the response to appear at (0.8, 0.8) after the bias adjustment. Residual error was calculated for all trials and then averaged within each participant. We also checked the correlation of the responses along the two axes, which was only r(11,512) = 0.016, confirming that it is appropriate to treat the two axes as independent.

The four parameters obtained by fitting (Bx, By, and WB related to the bias; the residual error, a measure of precision) were compared across conditions. Each was submitted to an ANOVA as above. There were no significant effects of being a boundary on any of the four parameters. 95% bootstrapping intervals on η^2^ reached 0.068 at the highest. In contrast, looking like a boundary had significant effects on the bias’s y-axis placement (By) and the bias’s weight (W_B_; Figure 5, Table 3). These effects were driven by the *cones* condition having a higher bias weight and placing the bias at a lower point on the y-axis (down in figures; see Figure 4). In summary, the pattern of responses was somewhat different for *cones* than the other conditions, biasing responses to a different place in the space and giving that bias some additional weight. This formally confirms that the differences in Figure 4 are statistically significant. However, we found no evidence that behaviour was different in *plywood* versus *pass through* or *pass over* – see also the 95% confidence intervals in Figure 5. This does not provide any evidence for an effect of looking like a boundary.

**Figure 5:**
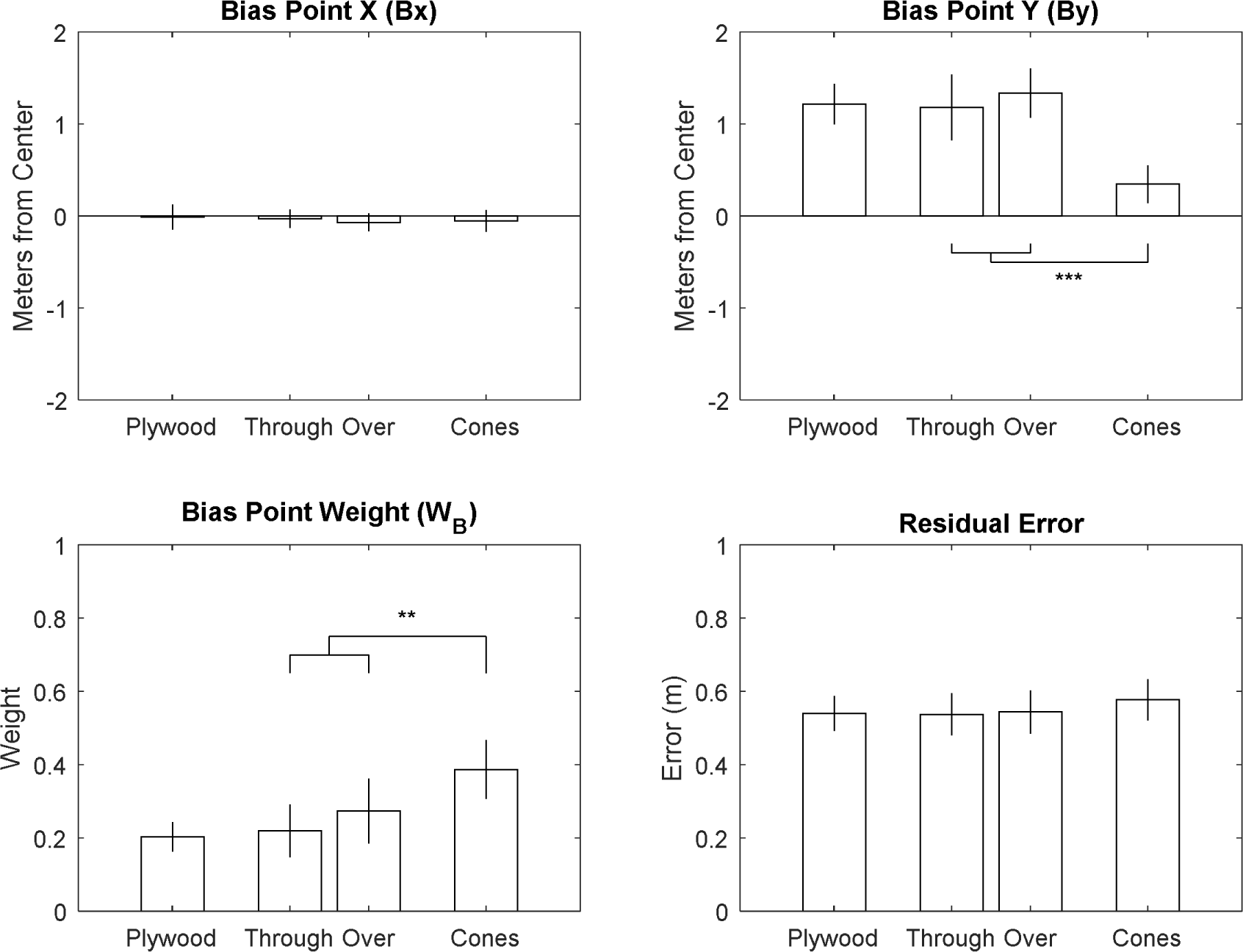
Average Fitted Values by Condition. There was a significant effect only on the bias point’s weight (W_B_) and placement along the y-axis (By). This was driven by differences between the *cones* condition (far right) and *pass through/over* (middle two). In other words, it is due to looking like a boundary but not due to being a (physical) boundary. Looking just at the three conditions with wall-like items, the 95% confidence intervals all overlap within each of the four outcomes. For example, a 20% bias weight is within the confidence intervals for *plywood, pass through*, and *pass over*. *** *p* < .001; ** *p* < .01

An alternative model-based version of this analysis was also performed, detailed in Appendix A. This analysis used Deviance Information Criterion (DIC), which can select simpler models over more complex ones on the basis of predicting future data accurately (Spiegelhalter, Best, Carlin, & van der Linde, 2002). In that analysis, a model with no difference between *plywood*, *pass through*, and *pass over* was preferred over one that includes a difference between *plywood* versus *pass over* and *pass through*. In other words, we did not just fail to find an effect of being a (physical) boundary, but found a positive reason to leave an effect of being a boundary out of the model.

A few additional results are not particularly useful for deciding among our hypotheses but still show that the task is a sensible measure of spatial cognition (Figure 6). For these, we worked with the raw error (the absolute distance between target and response; not the residual error as above). For each participant, an unstandardized beta coefficient was calculated with absolute rotation as the predictor variable and raw error as the outcome. The average beta value was significantly above zero, t(79) = 11.65, p < .001, mean of .0016 meters per degree. In other words, being displaced further led to higher error. This is in line with well-established advantages for spatial recall from similar versus different viewpoints (e.g. King, Burgess, Hartley, Vargha-Khadem, & O’Keefe, 2002). Average raw error was also lower for walking to the target than pointing to it, t(79) = 3.55, p < .001, Cohen’s d = 0.40, perhaps because moving gives access to additional viewpoints of the space. Average raw error also varied by block number, F(3, 319) = 6.66, p < .001, eta squared η^2^ = .06. This appears to just be an ordinary learning effect, with lower average error in later blocks.

**Figure 6:**
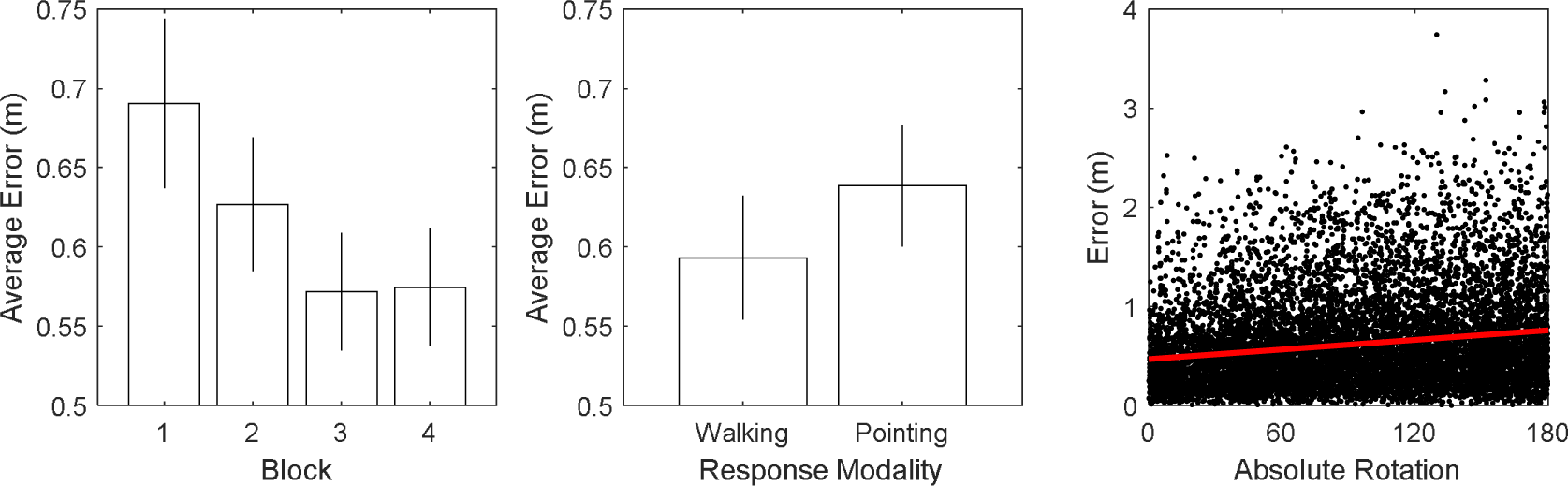
Additional Effects on Raw Error (without bias correction). Average raw error decreased as participants practiced the task. Walking to the targets was slightly easier than pointing to them, perhaps due to exposure to additional viewpoints. As expected, a smaller absolute rotation (i.e. a shorter displacement and smaller change in viewpoint) tended to result in lower raw error. All three main effects presented here are statistically significant at p < .001. While these effects do not specifically decide among our hypotheses, they do suggest that the task was functioning normally as a spatial cognition task.

## Discussion

Participants were given a typical spatial cognition task, indicating where a remembered target was from a new viewpoint. The design dissociated whether the environments contained physical boundaries and whether the environments had items that looked like boundaries (Figure 1; Table 1 and 2). We found zero significant effects of being a boundary across the five different analyses performed. The highest 95% confidence interval on η^2^ suggested that being a boundary could only explain up to 6.8% of the variance. However, we found two significant effects of looking like a boundary, which shifted the bias in the environment and gave the bias more weight. DIC results (in Appendix A) agree, suggesting that the best model excludes an effect of being a boundary. As a whole, this suggests that an item’s appearance is largely driving the observed effects when typical boundaries and non-boundaries lead to different patterns of responses in a spatial cognition task. When we compared experimental conditions that were visually identical, but where one condition contained an item that was a physical boundary and the other condition did not, confidence intervals were small and no effects were significant.

It might also be valuable to view this set of results from another perspective. In the usual design for an experiment on this topic, we would compare typical boundaries versus typical landmarks in terms of a spatial cognition measure. We included that comparison in a subset of the conditions used here. The *plywood* condition contained typical boundaries and the *cones* condition contained typical landmarks. When comparing those two conditions, we observed a clear difference in the outcome measure. This is in line with a variety of similar previous results (Doeller & Burgess, 2008; Keinath et al., 2017; Lee, 2017). In addition, we also added two control conditions (*pass over*/*through*). These conditions did not contain boundaries, like *cones*, but did have items that looked like boundaries, like *plywood*. Resulting behaviour was much more similar to *plywood* than *cones*. This emphasizes the importance of looking like a boundary over being a boundary. This also converges with a previous report on social boundaries, which were not found to affect spatial cognition differently than visually-identical non-boundaries (McNamara, 1986).

We no longer have to rely entirely on evidence from experiments with visual confounds to make conclusions about the effects of boundaries. We can now create and evaluate theory on the basis of observing what happens when two conditions are physically different but visually identical. The results fit a theory that focuses on the presence of visually extended 3D surfaces (Lee, 2017; Lee & Spelke, 2011), a visual aspect of looking like a boundary. Under this theory, all three conditions with walls visually presented a visually extended 3D surface, so it is expected that spatial coding would be the same. The cones lack such a surface, so it is expected that coding would differ from the other conditions.

A rational or Bayesian account of spatial cognition (Twyman, Holden, & Newcombe, 2018; Xu, Regier, & Newcombe, 2017) can also explain some critical features of the data. Biases in the *cones* condition shifted more towards the center of the target area. Responses near the center remove the possibility of making the largest possible errors (it is not possible to make an error larger than the radius of the target area). The traffic cones are likely to be more difficult to use for spatial encoding, and thus memory noise might be very high (Mou & Zhou, 2013), leading rationally to the bias towards the center to compensate. However, it is much less clear what rational purpose the bias towards the back wall would serve in the other conditions.

A domain-general account using learning theory could also explain the present data (Pearce, 2009). Under this kind of account, the three conditions with walls all provide the same visual features to associate a target with. No effect of different wall types is expected. In contrast, the cones provide fewer points through the space for association and might be less visually salient (Kosaki et al., 2013). Therefore, an effect of walls versus cones is expected.

These results are much more difficult to accommodate under a theory that places special focus on (physical) boundaries, disregarding how they look (Doeller & Burgess, 2008; Gallistel, 2017). Doeller and Burgess (2008) defined a boundary as an obstacle that subtends a large horizontal angle at the animal. They varied one of the visual confounds in their study (tripling the height of the landmarks in one condition), which did not affect the pattern of results. Gallistel (2017) defined a boundary as an object with an impact on what is navigable, without reference to visual features. Both concluded a special role for boundaries. Under these theories, we would expect the *plywood* boundaries here to be selectively involved in the formation of a ‘cognitive map’ in the hippocampus (Doeller et al., 2008), leading to large effects on how (or even if) locations are encoded (Gallistel, 2017). Instead, it fits the current data better to posit that both boundaries and non-boundaries alike have access to this system when they are visually identical.

While this helps identify the mechanism by which typical boundaries have special effects, it does not diminish how important these special effects can be in everyday life. In the last ten years, many articles have explored the neural mechanisms that rely heavily on typical boundaries to organize the way they represent space (Batty et al., 2010; Doeller et al., 2008; Ferrara & Park, 2016; Julian, Ryan, Hamilton, & Epstein, 2016; Keinath et al., 2017; Lee et al., 2018; Lescroart & Gallant, 2018; Lever et al., 2009). These patterns of coding and their special relation to typical boundaries are crucial to a full understanding of how humans and other mammals process the space around themselves.

### Potential Limitations

At first, ‘teleporting’ participants between encoding and recall may seem like a weakness of the present study, as this may damage their sense that the (physical) boundaries were real boundaries. However, participants did not view the testing room and did not attempt to touch the boundaries again. This means that they did not have any reason to believe the boundaries were not still real. For this to be an issue, disorientation itself would have to make participants think that nearby objects will have changed in terms of their ability to block navigation. Further, a disorientation procedure actually prompts the particular use of typical boundaries in recovering orientation and hippocampal coding (Cheng et al., 2013; Hermer & Spelke, 1994; Keinath et al., 2017; Lee, 2017). In addition, the lack of such a procedure would allow the task to be solved through dead reckoning without attending to any objects in the environment (McNaughton, Chen, & Markus, 1991). Taken together, this means that a ‘teleporting’ procedure is the most appropriate way to examine any potential effects of being a boundary.

Beyond the specific choice to virtually ‘teleport’ participants, there might also be more general concerns regarding the use of immersive virtual reality. This can be a weakness in some designs, but not the present study. The fidelity of immersive virtual reality simulations for studying spatial cognition has undergone extensive examination, far more than most Psychology methods, often uncovering systematic linear underestimation of egocentric distances (Renner, Velichkovsky, & Helmert, 2013). However, this effect is generally attenuated greatly, if not eliminated, by a “walking intervention” (i.e. short experience moving through the space via walking), which our participants had. It is also a much smaller concern for modern head-mounted displays (Kelly, Cherep, & Siegel, 2017) like the one used here. Several complex spatial effects have been shown to be the same in immersive virtual reality and real stimuli (Kelly, Avraamides, & Loomis, 2007; Kelly & McNamara, 2008; Negen, Heywood-Everett, Roome, & Nardini, 2018; Negen & Nardini, 2015; Williams, Narasimham, Westerman, Rieser, & Bodenheimer, 2007). In terms of boundaries specifically in immersive virtual reality, participants go to great effort to avoid striking things that look like boundaries even when participants know the items are not boundaries (Fink, Foo, & Warren, 2007), with their paths in novel virtual reality environments being very closely predicted (R^2^ > .95) by parameters gathered in dissimilar real environments. More troubling issues may be present when using desktop virtual reality instead of immersive virtual reality, yet even that approach captures individual differences that are also seen in the same participants in real environments (Richardson, Montello, & Hegarty, 1999). Perhaps even more to the point for the present study, the crucial difference between the *plywood* vs. the *pass through* and *pass over* conditions was not delivered through VR at all – it was a real difference between the subject’s real hand touching a piece of real plywood or not.

### Future Work

A large-scale study where several different visual item parameters are manipulated systematically might be an invaluable resource in clarifying the precise visual mechanisms that were studied here. For example, one might hypothesize that vertical extent is the reason why behaviour differed between *cones* and *pass through*, but this might make take several forms. There might be a strong discontinuity in the effects due to vertical extent (items may change semantically from a ‘line’ to a ‘kerb’). There might be a smooth function. The precise effects of different visual parameters could and should be characterized to gain a full model of how spatial memory is affected by different environments.

## Conclusion

Experiencing that an item is a (physical) boundary, independent of how much it looks like a typical boundary, does not directly affect adult participants’ coding of the locations around it. In contrast, non-boundaries that vary in how they look can lead to substantial differences in the pattern of responses. This is the clearest evidence to date that the typical boundaries in natural scenes have their particular effects on spatial cognition because of visual aspects such as horizontal extent, the presence of a visually extended 3D surface, or large-scale structure – not because they limit navigation.

## Supporting information

Raw Data

## Acknowledgements

Funding: This work was supported by grant 220020240 from the James S. McDonnell Foundation 21st Century Science Scholar in Understanding Human Cognition Program and grant ES/N01846X/1 from the Economic and Social Research Council of the United Kingdom. This project has received funding from the European Research Council (ERC) under the European Union’s Horizon 2020 research and innovation programme (grant agreement No. 820185).

## Appendix A

This appendix narrates and then details a DIC version of the main analysis. We modelled these data in a way where each person has a specific point that their responses are biased towards by a certain strength, corrupted by random memory noise. Based on Figure 6, the model also features a term for the rotation amount and the block number to influence precision. We can then use Deviance Information Criterion (Spiegelhalter et al., 2002) to assess how different nominal groupings of the different conditions affects the predictive value of the model (i.e. to test for/against differences between conditions). For example, DIC should improve when we collapse the *pass over* and *pass through* conditions if their relevant parameters are actually quite similar, since it means that more data can be used to predict future participants in either condition. In contrast, DIC should worsen if they are actually dissimilar, since it would then be more accurate to just use each condition’s separate data to predict future participants.

### Model Analyses

The description of our formal model-based analysis proceeds in four stages. First, we describe the model (a typical model of Bayesian reasoning with a prior and likelihood using bivariate normal distributions and additive precision) in a way that is designed to help the reader gain an intuition of how it functions. Second, we state the main result. Third, we look at the posterior distributions of the model to see what is driving the main result. Fourth, full formal details are provided.

#### Model description

After a target is shown (Figure A1, left panel) and removed, its memory trace degrades during the ‘teleportation’. By the time recall is occurring, the memory trace (Figure A1, middle panel) is a draw from a bivariate Gaussian with the target as the mean. In addition to the trace, there is a prior distribution that the participant is using. With their relative precisions used as weights, the prior’s center and the memory trace are averaged to determine the response (Figure A1, right panel).

**Figure A1.**
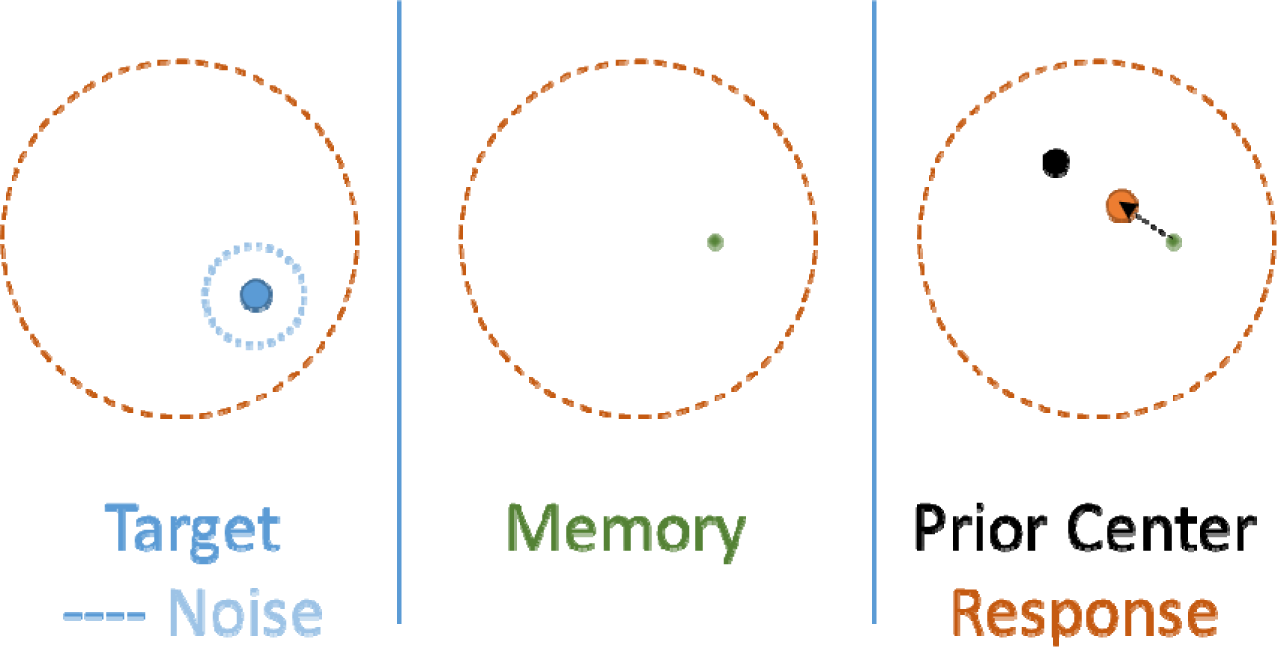
An example trial of how the model works. The memory trace of the target in the left panel degrades over the ‘teleport’ to the place in the middle panel. The participant then takes a weighted average of their memory (green dot) and their subjective prior (black dot) to determine the response (orange dot) in the right panel.

This model therefore has four parameters per participant: the prior’s center on the x-axis (*Prior X*), the prior’s center on the z-axis (“forwards” for the participant standing in the encoding circle; *Prior Z*), the precision of the memory traces (*Memory Tau*), and the precision of the prior distribution (*Prior Tau*). The bias they show is determined by the prior’s center (where they show a bias towards) and by the prior precision (how strong that bias is). Their memory precision is generally the precision of interest, controlled directly by Memory Tau, but the overall precision of their responses is additively determined by the memory precision and the prior precision.

It is worth noting that the mathematical mechanics involved in this model can be re-framed equally as a bias existing towards a certain point with a certain strength, trading off against their memory and their confidence in that memory. It could also possibly be framed as over- or under-representation of equal portions of the space. We do not have the data to verify the extensive predictions that come along with assuming fully optimal Bayesian reasoning.

The rest of the model then assumes that individuals, with their four individual-level parameters, are drawn from a condition-level distribution with two parameters per individual-level parameter: one for the mean value within the condition (Prior X Mean, Prior Z Mean, Memory Tau Mean, Prior Tau Mean), and another for the spread (Prior X Tau, Prior Z Tau) or the shape/spread of the Gamma distribution that they are drawn from (Memory Tau Alpha, and Prior Tau Alpha). The posterior fitting process flows in both directions, with the condition-level distributions responding to individual-level parameters and vice versa.

We also created a version to look for interaction effects. In this version, each participant has a separate Prior X and Prior Z parameter for walking and pointing, but they are correlated in a bivariate normal. The Memory Tau and Prior Tau parameters are replaced by their logarithms, which are given the same treatment: separate but correlated across modalities.

#### Main result

The main result is that the Deviance Information Criterion (DIC; see Table A1) is best when grouping the conditions with walls together into one nominal condition with the same group-level parameters and keeping the *cones* condition separate, while also keeping the pointing and walking trials together within conditions (second row to the bottom, highlighted). This is the best DIC by a margin of 30 (49 below baseline model, whereas the next best is 19 below). For calibration, imagine two models with the same number of effective parameters. If one fits the data 150 times worse on average, that one will have a mean deviance and thus DIC that is higher by 10. DIC also has an explicit penalty for model complexity in units of deviance. The selected model has the lowest (best) DIC for a combination of both reasons, having the best average fit (by 11) and the second-smallest complexity penalty (smallest was the model with no effect of condition whatsoever).

**Table A1.** DIC Results.

| Condition Groupings | Interpretation | Mean Deviance | p<sub>v</sub> | DIC |
| --- | --- | --- | --- | --- |
| 1 = Plywood,<br>2 = Pass Through,<br>3 = Pass Over,<br>4 = Cones | (Full Effects Baseline)<br>Every condition difference matters | 28,051 | 427 | 28,478<br>(+/-0) |
| 1 = {Plywood, Pass Through, Pass Over, Cones} | (Null Effects) No condition difference matters | 28,133 | 391 | 28,523<br>(+45) |
| 1 = Plywood,<br>2 = {Pass Through, Pass Over},<br>3 = Cones | Both boundary status and visual covariates matter | 28,052 | 422 | 28,474<br>(-4) |
| 1 = Plywood,<br>2 = {Pass Through, Pass Over, Cones} | Boundary status matters, but not visual covariates | 28,044 | 415 | 28,459<br>(-19) |
| 1 = {Plywood, Pass Through, Pass Over}<br>2 = Cones | Visual covariates matter, but not boundary status | 28,033 | 396 | 28,429<br>(-49) |
| 1 = {Plywood, Pass Through, Pass Over, Cones} but with separate correlated parameters for walking and pointing | Separation of parameters by response modality (potential interaction effects)* | 32,189 | 3,147,591 | 3,179,779<br>(+3x10<sup>6</sup>) |

\*Note: for brevity, other groupings are omitted for the interaction effects, but they were all worse by margins are that functionally infinite (mean deviance up by >6,000, total DIC up by millions).

This favors an interpretation that visual covariates matter, but not true boundary status. More formally, it suggests that the best way to predict future data is to use all the recorded participants from the *plywood, pass over* and *pass through* conditions together to predict the performance of a new participant in any of those three conditions, but to only use the recorded *cones* data to predict the performance of a new participant in the *cones* condition. Note that this is the version where walking and pointing are kept together; all models that separated these suffered an enormous complexity penalty for nearly doubling the number of nominal parameters, but did not gain anything notable in terms of mean deviance.

#### Posterior distributions

What is driving the differences in DIC? An examination of the posterior distributions (Figure A2, *cones* in purple) using the full effects baseline model shows three places where the *cones* condition diverges from the rest to a notable degree: the average center of the inferred prior on the z axis is lower for the *cones* condition than the other conditions, the mean precision of the inferred prior is higher, and the mean precision of their memory is lower. In other words, on average, the biases they showed were pulled towards a different place (nearer the encoding point), and pulled there stronger, plus they had lower precision (more noise) in their memory of locations in the task.

**Figure A2.**
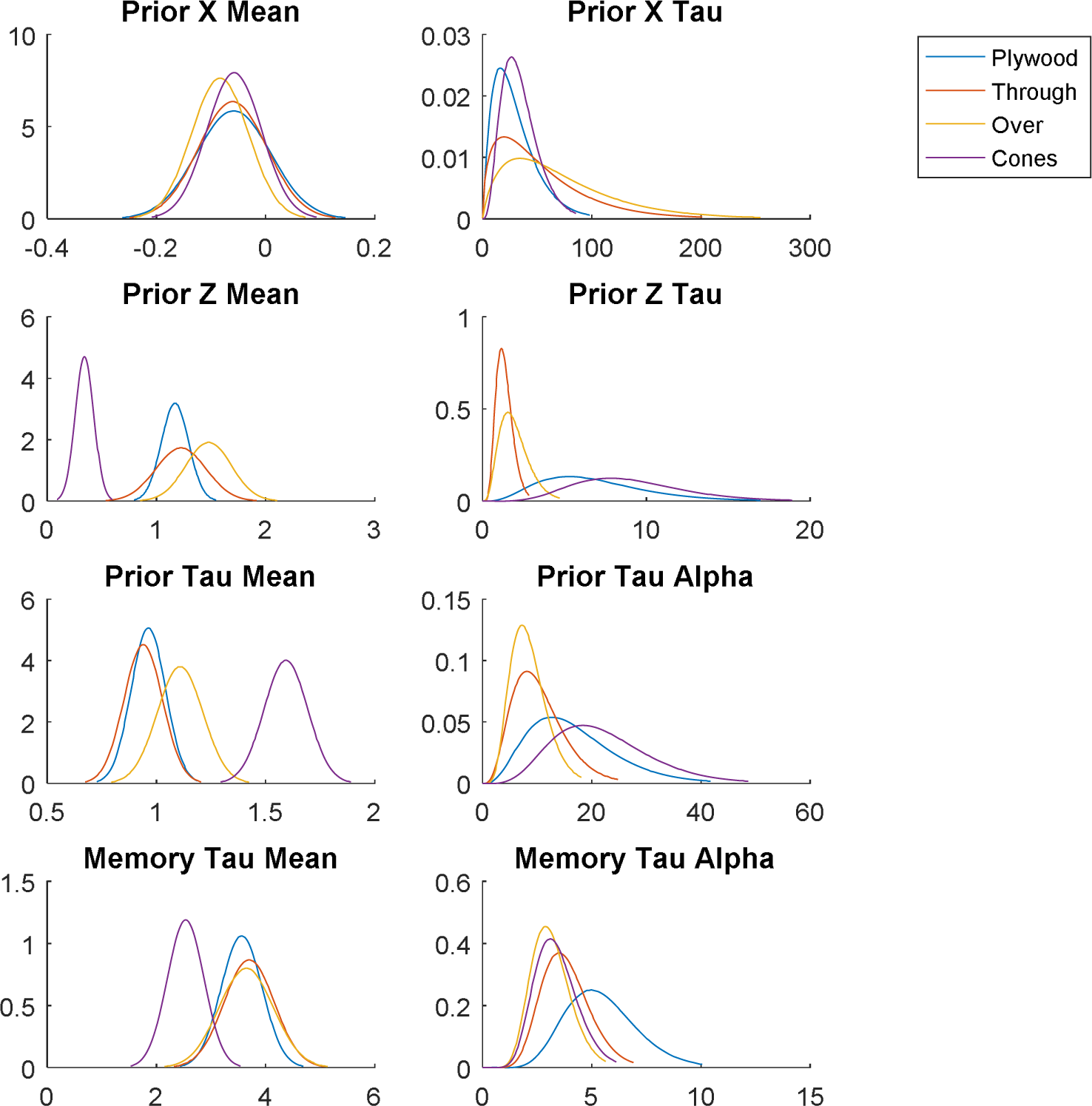
Posterior distributions for the four different conditions over the eight condition-level parameters. The differences in the *cones* (purple) condition explain how that condition differed and why DIC was lowest when grouping the others together. All units are in meters.

To make the bias difference easier to visualize, we also plot a 95% credible region around the mean prior point for each condition in Figure A3. Compare with the descriptive Figure 4 above to see that these fits are sensible. This is interesting because the *cones* fit is both quantitatively and qualitatively different. Since it is near the center of the actual area where the targets appear, it might reflect a default strategy of pointing near the center to minimize error when memory is not trustworthy.

**Figure A3.**
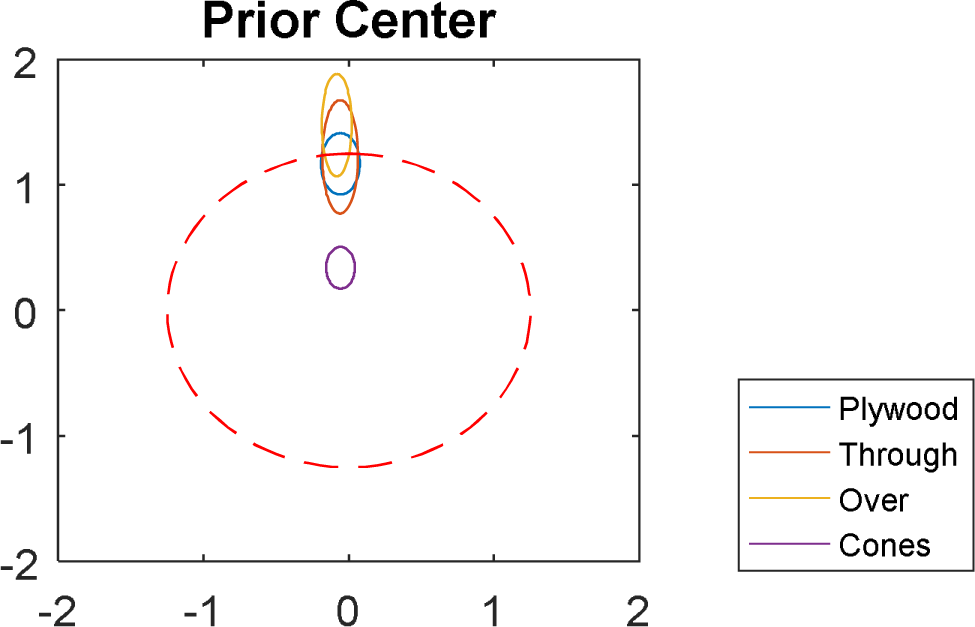
Visualization of the different prior centers on average over participants in each condition. Each ellipse represents a 95% credible region.

In contrast to the *cones*, the *plywood* condition is not clearly separated from the other two conditions with walls on any condition-level parameter. It is perhaps closest to showing a difference with the *pass through* and *pass over* conditions in terms of the Prior Z Tau parameter, which is somewhat awkward to interpret even if we ignore the overall DIC results; it would mean that participants in that condition were more consistent in where they placed the center of the prior on the z axis (but not the x axis, and arriving at the same mean placement).

For completeness, we also report that the mean posterior Rotation Beta was -.006 (95% Credible Interval: -.0064 to -.0056), suggesting that a trial where participants were rotated to the far side had 2.94 times the precision (about 58% of the standard deviation) of trials where they did not rotate at all. The effects of the blocks, in order, were 0 (fixed), 0.22 (0.16 to 0.28), 0.40 (0.34 to 0.46), and 0.38 (0.33 to 0.44). For reference, a value of 0.40 translates to 49% more precision (82% of the standard deviation).

### Formal details

This section gives a full formal description of the model that was applied to the data.

The model has 4 global parameters, 8 condition-level parameters per condition, and 4 individual-level parameters per participant.

- Rotation_Beta (global) controls how much rotating a participant (i.e. how far away the recall viewpoint was from the encoding viewpoint) affects the precision of their memory. It was given a prior of a normal distribution with mean 0 and precision 0.01.
- Block_Beta[2-4] (global) control how much a trial existing in blocks 2-4 affects the precision of memory. They were given priors of normal distributions with mean 0 and precision 0.01. A trial in block 1 has this set to 0, which does not affect precision, as a way of identifying the model.
- Prior_X_Mean (condition) is the center of the condition-level distribution of the priors’ x-axis values. It was given a normal prior with mean 0 and precision 0.01.
- Prior_X_Tau (condition) is the precision of the condition-level distribution of the priors’ x-axis values. It was given an exponential prior with rate of 0.01.
- Prior_Z_Mean and Prior_Z_Tau (condition) are the same as the X-axis parameters above, except on the Z axis.
- Memory_Tau_Alpha (condition) controls the shape of the gamma distribution from which individual memory precisions are drawn for participants in that condition. It was given an exponential prior with a rate of 0.01.
- Memory_Tau_Beta (condition) controls the rate of the gamma distribution from which individual memory precisions are drawn for participants in that condition. It also has a prior distribution of an exponential with rate of 0.01.
- Prior_Tau_Alpha and Prior_Tau_Beta (condition) are the same as just above but for the precision of the prior for that participant in that condition.
- Prior_X and Prior_Z (individual) together encode the center point of each participant’s inferred prior. Prior_X is drawn from Normal(Prior_X_Mean, Prior_X_Tau ^−1/2^) and the same form for the Prior_Z.
- Memory_Tau and Prior_Tau (individual) together encode the precision of the participant’s memory for a trial with 90 degree absolute rotation in block 1, and the precision of the inferred prior. Memory_tau is drawn from a gamma distribution with shape Memory_Tau_Alpha and rate Memory_Tau_Beta, and the same form for Prior_Tau.

For a given pair of target and response, we then assume that a response is drawn from a bivariate normal with calculated (non-stochastic) parameters for the means and standard deviation, assuming the covariances to be zero and the standard deviation to be the same in both axes.

The standard deviation is the total precision to the power of negative one half. The total precision has two additive components. The first is the prior precision. The second is the memory precision, which is Memory_Tau for that participant times exp(Rotation_Beta*abs(90-Rotation)) times exp(Block_Beta[Block]).

The mean is a weighted average between the target’s location and the prior’s center. The weight for the target’s location is calculated by dividing the memory precision by the total precision. The weight for the prior’s center is calculated by dividing the prior precision by the total precision. This is done in both axes.

This model purposefully makes several omissions that we can now explain. There is no explicit correlation between responses within the same trial. This was driven empirically, as Pearson’s r values between the (response-target) values were relatively low within the same trial, between −0.05 and 0.25. We also saw that the model’s assumption structure provide certain kinds of similarity between responses on the same trial anyway, such as a common total precision, as they all share the same parameters.

The model also omits different effects of block or rotation by condition. This was done for two reasons. First, because we wanted to have a model where the response modalities were separated into different conditions. Having these effects vary by condition would then become much less stable, as participants only completed two block numbers in each response modality. Second, we did not have any a priori reason to think they would vary by condition.

The interaction model replaced Memory_Tau and Prior_Tau with the logarithms, then made each participant-level parameter subject to a bivariate normal distribution separated by response modality. The priors were a conjugate normal-wishart model: means of zero for the means, precisions of .01 for the means, zero correlation in the priors over means, and a wishart distribution with a rank of 2 and an identity matrix for omega.

Below is the precise WinBUGS Code used for the main model:

~~~
model{
for (i in 1:nConditions){
  CenterXMean[i] ∼ dnorm(0,.01)
  CenterXTau[i] ∼ dexp(.01)
  CenterZMean[i] ∼ dnorm(0,.01)
  CenterZTau[i] ∼ dexp(.01)
  TauAlpha[i] ∼ dexp(.01)
  TauBeta[i] ∼ dexp(.01)
  TauMean[i] <- TauAlpha[i] / TauBeta[i]
  PriorTauAlpha[i] ∼ dexp(.01)
  PriorTauBeta[i] ∼ dexp(.01)
  PriorTauMean[i] <- PriorTauAlpha[i] / PriorTauBeta[i]
}
TauBlockEffect1 <- 0
TauBlockEffect2 ∼ dnorm(0,.01)
TauBlockEffect3 ∼ dnorm(0,.01)
TauBlockEffect4 ∼ dnorm(0,.01)
TauBlockEffect[1] <- TauBlockEffect1
TauBlockEffect[2] <- TauBlockEffect2
TauBlockEffect[3] <- TauBlockEffect3
TauBlockEffect[4] <- TauBlockEffect4
TauRotationEffect ∼ dnorm(0,.01)
for (i in 1:80){
  CenterX[i] ∼ dnorm(CenterXMean[ConditionByParticipant[i]],
      CenterXTau[ConditionByParticipant[i]])
  CenterZ[i] ∼ dnorm(CenterZMean[ConditionByParticipant[i]],
      CenterZTau[ConditionByParticipant[i]])
  Tau[i] ∼ dgamma(TauAlpha[ConditionByParticipant[i]],
      TauBeta[ConditionByParticipant[i]])
  PriorTau[i] ∼
      dgamma(PriorTauAlpha[ConditionByParticipant[i]],PriorTauBeta[ConditionByParticipant[i]])
}
for (i in 1:N){
  TauTrialPerception[i] <- Tau[Participant[i]] * exp(TauBlockEffect[Block[i]]) *
      exp(TauRotationEffect * Rotation[i])
  Weight[i,1] <- TauTrialPerception[i] / (TauTrialPerception[i] +
      PriorTau[Participant[i]])
  Weight[i,2] <- PriorTau[Participant[i]] / (TauTrialPerception[i] +
      PriorTau[Participant[i]])
  muX[i] <- Weight[i,1] * XTar[i] + Weight[i,2] * CenterX[Participant[i]]
  muZ[i] <- Weight[i,1] * ZTar[i] + Weight[i,2] * CenterZ[Participant[i]]
  TauTrial[i] <- TauTrialPerception[i]+PriorTau[Participant[i]]
  XRes[i] ∼ dnorm(muX[i],TauTrial[i])
  ZRes[i] ∼ dnorm(muZ[i],TauTrial[i])
}
}
~~~

